# A systematic review of MERS-CoV (Middle East Respiratory Syndrome Coronavirus) seroprevalence and viral RNA prevalence in dromedary camels: implications for animal vaccination

**DOI:** 10.1101/574103

**Authors:** Amy Dighe, Thibaut Jombart, Maria D. Van Kerkhove, Neil Ferguson

## Abstract

Human infection with Middle East Respiratory Syndrome Coronavirus (MERS-CoV) is driven by recurring dromedary-to-human spill-over events, leading decision-makers to consider dromedary vaccination. Dromedary vaccine candidates in the development pipeline are showing hopeful results, but gaps in our understanding of the epidemiology of MERS-CoV in dromedaries must be addressed to design and evaluate potential vaccination strategies. We systematically reviewed the published literature reporting seroprevalence and/or prevalence of active MERS-CoV infection in dromedary populations from both cross-sectional and longitudinal studies, including 60 studies in our qualitative syntheses. MERS-CoV seroprevalence increased with age up to 80-100% in adult dromedaries supporting geographically wide spread endemicity of MERS-CoV in dromedaries in both the Arabian Peninsula and countries exporting dromedaries from Africa. The high prevalence of active infection measured in juveniles and at sites where dromedary populations mix should guide further investigation – particularly of dromedary movement – and inform vaccination strategy design.

## Introduction

Since the first human case of Middle East Respiratory Syndrome Coronavirus (MERS-CoV) infection was detected in 2012 (1), a substantial evidence base has built up showing dromedary camels to be the zoonotic source of this virus (2). MERS-CoV circulates extensively in dromedary populations causing no impactful disease. Human infection, however, is associated with a measured case fatality ratio of around 35% (3). Following spillover events, human-to-human transmission of MERS-CoV is relatively inefficient and limited to close, unprotected contact environments such as hospitals (4, 5). Phylogenetic analysis of viral sequences isolated from dromedaries and humans indicates that hundreds of camel-to-human spillover events are likely to have occurred since 2012 (6). Taken together, recurring dromedary-to-human transmission is driving ongoing human infection.

The key role of dromedaries in human MERS-CoV infection has led decision-makers to consider dromedary vaccination as part of MERS-CoV prevention interventions (2). Dromedary-targeted vaccine candidates in the development pipeline are showing promising results and include an orthopox-virus based vaccine capable of greatly reducing viral shedding in dromedary challenge studies (7).

However, vaccine strategy evaluation is currently precluded by gaps in the understanding of the epidemiology of MERS-CoV in dromedaries. The dromedary population is highly heterogeneous and spans a wide geographic area stretching from West Africa through to the Middle East and parts of Asia. Knowing when and where dromedaries would need to be targeted, and the likely impact of vaccination, is necessary before further consideration of dromedary vaccination in the wider socioeconomic and cultural context.

Here, we systematically reviewed published studies that measured MERS-CoV antibody seroprevalence in dromedaries and/or prevalence of viral RNA in dromedaries. Assuming assay specificity and long-term presence of antibodies after infection, seroprevalence can be used to estimate what proportion of a dromedary population has ever been infected with MERS-CoV. Broken down by age class, this can tell us when most animals encounter the infection for the first time. Additionally, although whole-virus isolation and culture is necessary to confirm whether the shedding is infectious, detection of viral RNA through RT-PCR can be used as a proxy for the prevalence and distribution of infectious dromedaries (8–10).

By conducting a qualitative synthesis of the study findings, considering reported heterogeneities, and summarising the results of longitudinal studies of infection and immunity, we aim to assess the extent of current understanding of MERS-CoV epidemiology in dromedaries, implications for control, and gaps to be addressed going forward. We note that a similar systematic review of the literature up until May 2018 was unknowingly carried out in parallel to our own (11), with no discussion between the two groups. Here, we confirm and update the results of the parallel review, discussing our results in the context of potential animal vaccination and mathematical modelling of MERS-CoV in dromedary camels.

## Methods

We conducted a systematic review of studies published prior to 31st December 2018 reporting measures of seroprevalence or prevalence of MERS-CoV RNA in dromedary populations by searching EMBASE (12), MEDLINE (13) and Web of Science (14) using the search strategy in Fig 1. For a full list of search terms used and corresponding PRISMA flowchart see S1 Appendix and S2 Appendix.

**Figure 1.**
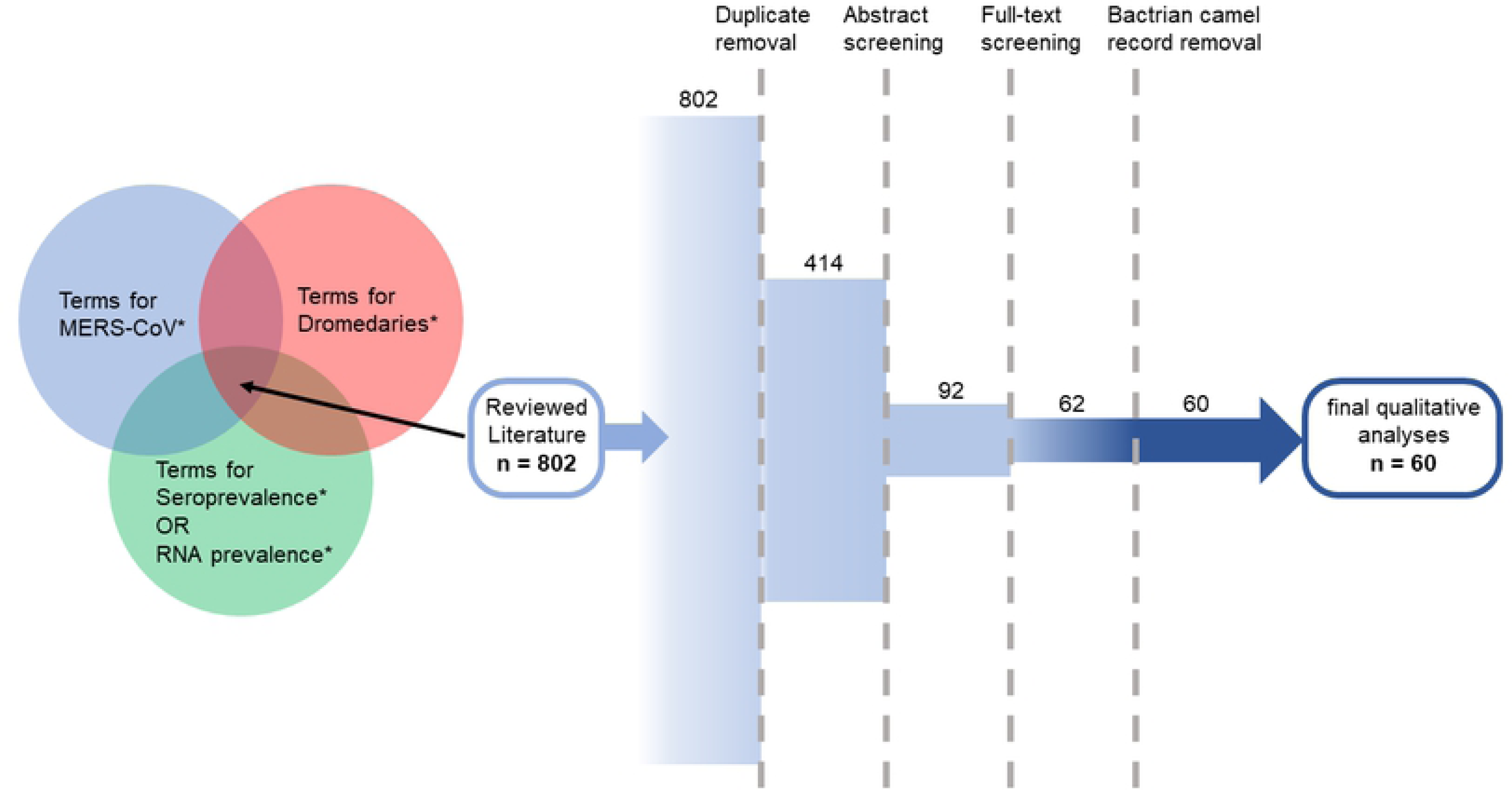
Published studies found with all three of our selected search term groups were then assessed against the exclusion criteria resulting in a final selection of 60 publications.

Records were excluded if they met the following criteria established prior to the search: opinion pieces, or reviews reporting no new data, studies investigating an aspect of MERS-CoV or another pathogen that did not involve dromedary samples or involved experimentally infecting dromedaries with MERS-CoV, and studies not available in English. Remaining studies were categorised as longitudinal or cross-sectional for qualitative synthesis.

When available, we took the results of neutralisation-based testing over methods that determined seropositivity based on antibody screening tests alone. Neutralising antibody tests are more specific and are the WHO (World Health Organisation) recommended method for confirming MERS-CoV seropositivity (9).

We extracted RNA prevalence values determined by RT-PCR (reverse-transcription polymerase chain reaction) of nasal swabs, ignoring any additional samples taken. Use of RT-PCR to test for the presence of at least two of the established genomic regions unique to MERS-CoV is the WHO and OIE (World Organisation for Animal Health) standard for detection of active MERS-CoV infection in dromedaries (9, 10), and viral RNA is most frequently and abundantly present in nasal swabs compared to other non-invasive samples (15).

Throughout this review, we use ‘calf’ to refer to animals under one-year-old, ‘juvenile’ ≤2 years old, and ‘adult’ ≥3 years old.

## Results

Our search retrieved 802 publications. Duplicates were detected and removed in EndNote X8.2 (16), leaving 414 unique publications. A further 322 records were excluded during abstract screening using the criteria given above. Of the remaining 92, full text screening proved that a further 30 records met the exclusion criteria. Two records sampled Bactrian camels (largely restricted to Central Asia, they have not yet been found to have been infected with MERS-CoV) (17–19), leaving 60 records pertaining to MERS-CoV seropositivity and/or RNA positivity in dromedaries (Fig 1). 55 of these described cross-sectional studies of dromedary populations, sampling each animal at a single time-point only (40 measured seroprevalence and 32 measured RNA prevalence). Note that 6 studies were designed to investigate groups of dromedaries that had been epidemiologically linked to human cases of MERS-CoV infection rather than conducting a systematic/random survey. Longitudinal studies that measured seropositivity and viral RNA shedding in the same animals at multiple time-points, featured in 11 publications.

### Seroprevalence – cross-sectional studies

Variously, studies conducted dedicated MERS-CoV sero-surveys, opportunistically tested samples taken for other means, tested stored sera or sampled dromedaries during human outbreak investigations. Not all studies used neutralisation-based testing to determine or confirm seropositivity, and, between those that did, cut-off titres for positivity varied (Table 1).

**Table 1:**
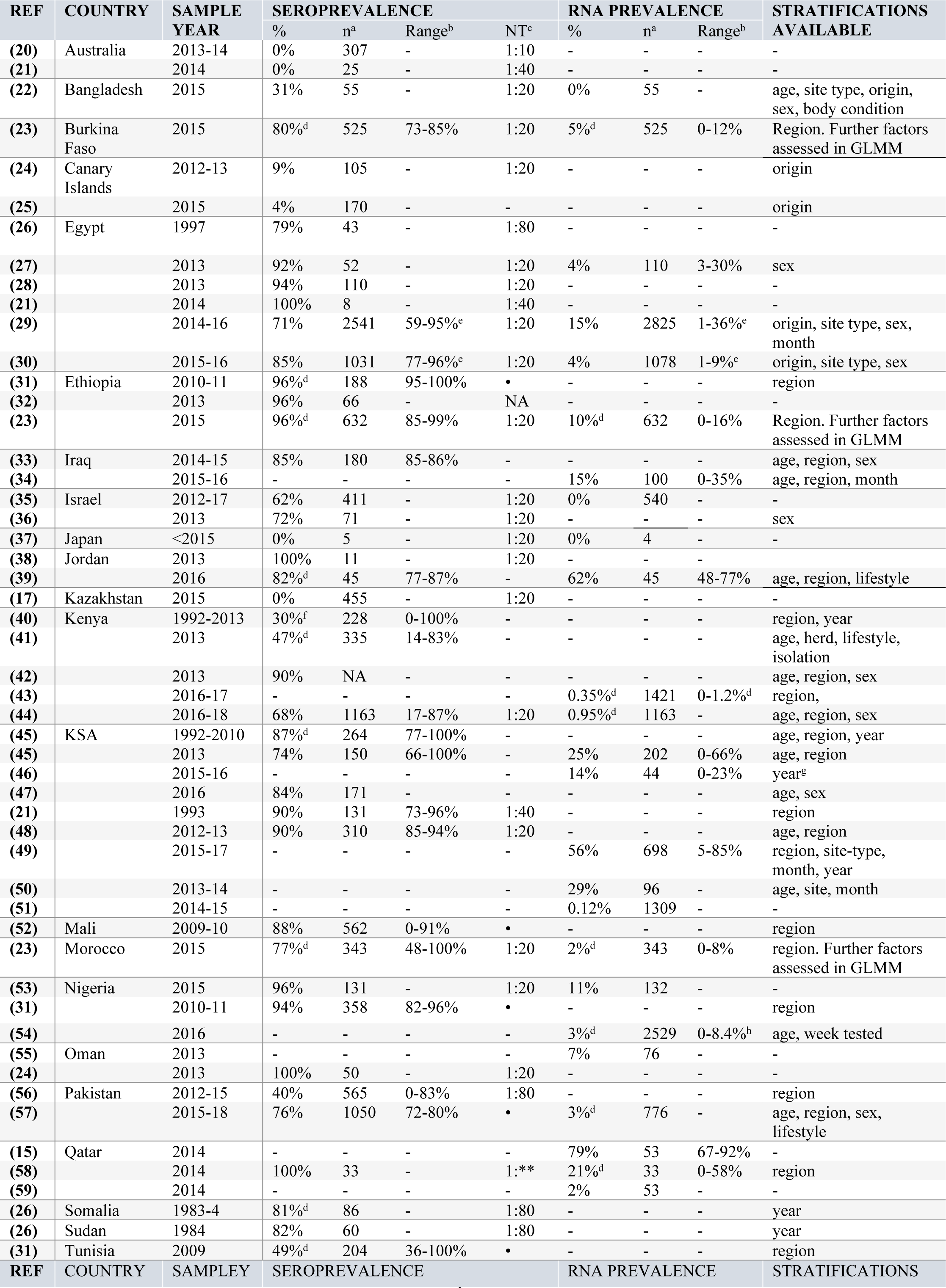
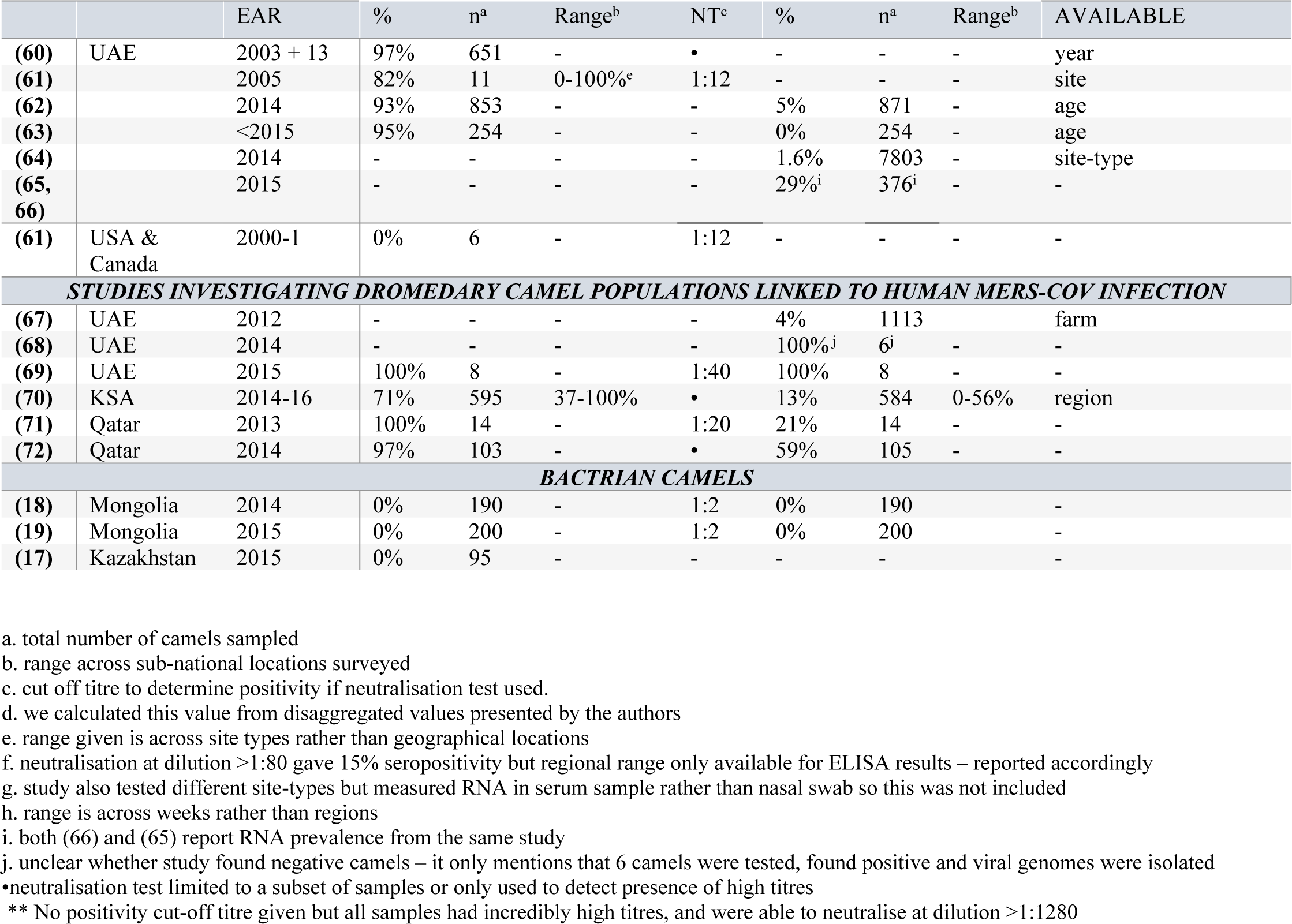
Cross-sectional surveys of MERS-CoV seroprevalence and RNA prevalence in camels

Seropositive dromedaries have been found in 20 of the 24 countries studied. See Table 1 and Fig 2 for a geographical map of seropositivity. MERS-CoV seroprevalence was between 71-100% in most country-level, age-aggregated study populations across West, North and East Africa and the Middle East. Exceptions included one of the two studies in Israel (62%), and 4-49% in Bangladesh, the Canary Islands, and Tunisia as well as one of the two studies in Pakistan and two of the four studies in Kenya. Samples taken from the large feral camel population in Australia were seronegative, along with dromedaries in Kazakhstan and in zoos in Japan, and Northern America (17, 20, 21, 24, 37, 61). No other species tested alongside dromedaries had neutralising antibodies except a small number of alpacas and llamas living in close quarters with dromedaries in Israel (35), and 1 sheep in Egypt (30).

**Figure 2.**
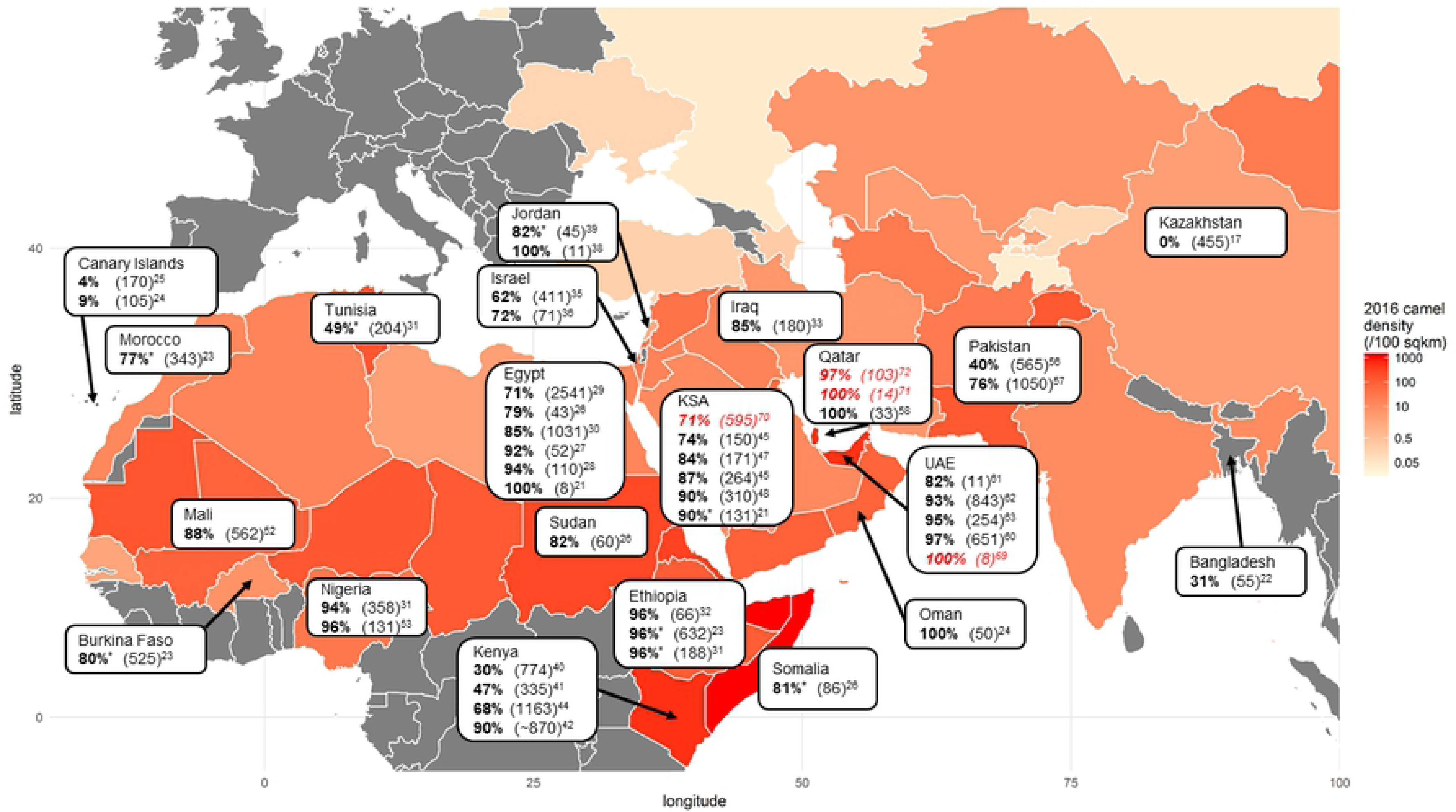
Measures of MERS-CoV seroprevalence in dromedaries, aggregated at the country level. Total sample size tested is given in parenthesis. Camel density is calculated using FAOSTAT country-level camel population data (73) and World Bank data on country surface area (74) (both for 2016). *value calculated by us from disaggregated sub-national measures of seroprevalence. Underlined italicised text highlights studies conducted in dromedary populations in response to an epidemiologically linked human MERS-CoV infection.

#### Factors effecting seroprevalence

##### Age

MERS-CoV seroprevalence was found to increase with dromedary age in 13 studies (Table 2, Fig 3). Although seroprevalence was 80-100% in adult dromedaries in most populations across the Middle East and the horn of Africa, juveniles in the same populations were repeatedly found to have lower and more variable seroprevalence (∼40-90%). One study disaggregated calf age through the first year of life. In this setting (UAE) calves had high seroprevalence increasing with age to 90% in 7-12-month-olds (63).

**Table 2:** Studies reporting seroprevalence stratified by age of dromedary

| COUNTRY | STRATIFIED SEROPREVALENCE | REPORTED TREND | REPORTED SIGNIFICANCE | REF |
| --- | --- | --- | --- | --- |
| BANGLADESH | <2yrs **9%** (n = 11)<br>≥2yrs **36%** (n = 44) | Higher in camels >2yrs | Not significant | (22) |
| BURKINA FASO,<br>ETHIOPIA AND<br>MOROCCO | Generalised Linear Mixed Model (n = 1500) | Increase with age | **p = 0.032** | (23) |
| EGYPT | <2yrs **52%** (n = 81)<br>≥2yrs **87%** (n = 950) | Higher in camels >2yrs | **p < 0.0001** | (30) |
| ETHIOPIA | 1- ≤2yrs **93%** (n = 31)<br>2-13yrs **97%** (n = 157) | None | Not significant | (31) |
| IRAQ | <2yrs **89%** (n = 44)<br>>2yrs **84%** (n = 136)<br>2-4yrs 81% (n = 58)<br>>4yrs 86% (n = 78) | Lower in camels 2-4yrs compared with <2yrs | Not significant | (33) |
| JORDAN | ≤2yrs **74%**<sup>a</sup> (n = 31<sup>a</sup>)<br>>2yrs **100%**<sup>a</sup> (n = 14<sup>a</sup>) | ELISA ratio higher in camels >3yrs | Significant p = NA | (39) |
| KENYA | <2yrs **29%**<sup>a</sup> (n = 141<sup>a</sup>)<br>>2yrs **61%** (n = 194)<br><6m 39% (n = 61)<br>6m-2yrs 21% (n = 80) | Higher in camels >2yrs than <6m | **p < 0.05** | (41) |
| KENYA | 1-4yrs 73% (n = 285)<br>4-6yrs 98% (n = 116)<br>6yrs 98% (n = 476) | Higher in camels >4yrs | **p < 0.05** | (42) |
| KENYA | <4yrs 36% (n = 319)<br>>4<7yrs 59% (n = 70)<br>>7yrs 82% (n = 760) | Higher in camels >7yrs | **p < 0.001** | (44) |
| KSA | ≤2yrs **55%** (n = 104)<br>>2yrs **95%** (n = 98) | Higher in camels >2yrs | **p < 0.0001** | (45) |
| KSA | ≤2yrs **73%** (n = 77)<br>>2yrs **93%** (n = 187) | Higher in older animals | Not presented | (45) |
| KSA | 1-2yrs **93%** (n = 71)<br>3-5yrs **78%** (n = 100) | Lower in camels >2yrs | **p = 0.03** | (47) |
| KSA | <1yr **72%** (n = 65)<br>>1yr **95%**<sup>a</sup> (n = 245)<br>1-3yrs 95% (n = 106)<br>4-5yrs 97% (n = 76)<br>>5yrs 92% (n = 63) | Higher in camels >1yr | **p < 0.01** | (48) |
| MALI | <2yrs **83%** (n = NA<sup>b</sup>)<br>3-8yrs 91% (n = NA<sup>b</sup>)<br>9-16yrs 88% (n = NA<sup>b</sup>) | None | Not significant | (52) |
| PAKISTAN | ≤3yrs 58% (n = 177)<br>3.1-10yrs 79% (n = 712)<br>>10yrs 81% (n = 161) | Lower in animals ≤3yrs | **p < 0.001** | (57) |
| PAKISTAN | ≤2yrs (n = 26/89)<br>2.1-5yrs (n = 62/208)<br>5.1-10yrs (n = 92/180)<br>>10yrs (n = 43/88) | Higher in older animals | **p < 0.001** | (56) |
| UAE | <1yr **85%** (n = 108) | Lower in calves <1yr | **p < 0.05** | (62) |
|  | >1yr **96%<sup>a</sup>** (n = 650)<br>2-4yrs 97% (n = 340)<br>>4yrs 96% (n = 310) |  |  |  |
| **UAE** | <1yrs **84%** (n = 121)<br>>1yrs **99%** (n = 133)<br>0-3m 75% n = 32)<br>4m 79% (n = 14)<br>5-6m 89% (n = 46)<br>7-12 90% (n = 29) | Increase with age | Not tested | (63) |
| **STUDIES INVESTIGATING DROMEDARY CAMEL POPULATIONS LINKED TO HUMAN MERS-COV INFECTION** |  |  |  |  |
| **KSA** | ≤2yrs **58%** (n = 25)<br>>2yrs **81%** (n = 344)<br>2.1-4yrs 77%<sup>c</sup> (n = 156)<br>4.1-6yrs 81%<sup>c</sup> (n = 98)<br>>6yrs 87%<sup>c</sup> (n = 90) | Higher in camels >2yrs | **p = 0.003** | (70) |

- a. age stratified results were calculated by us, using disaggregated results presented by authors. - b. number of animals in each age class not supplied

**Figure 3.**
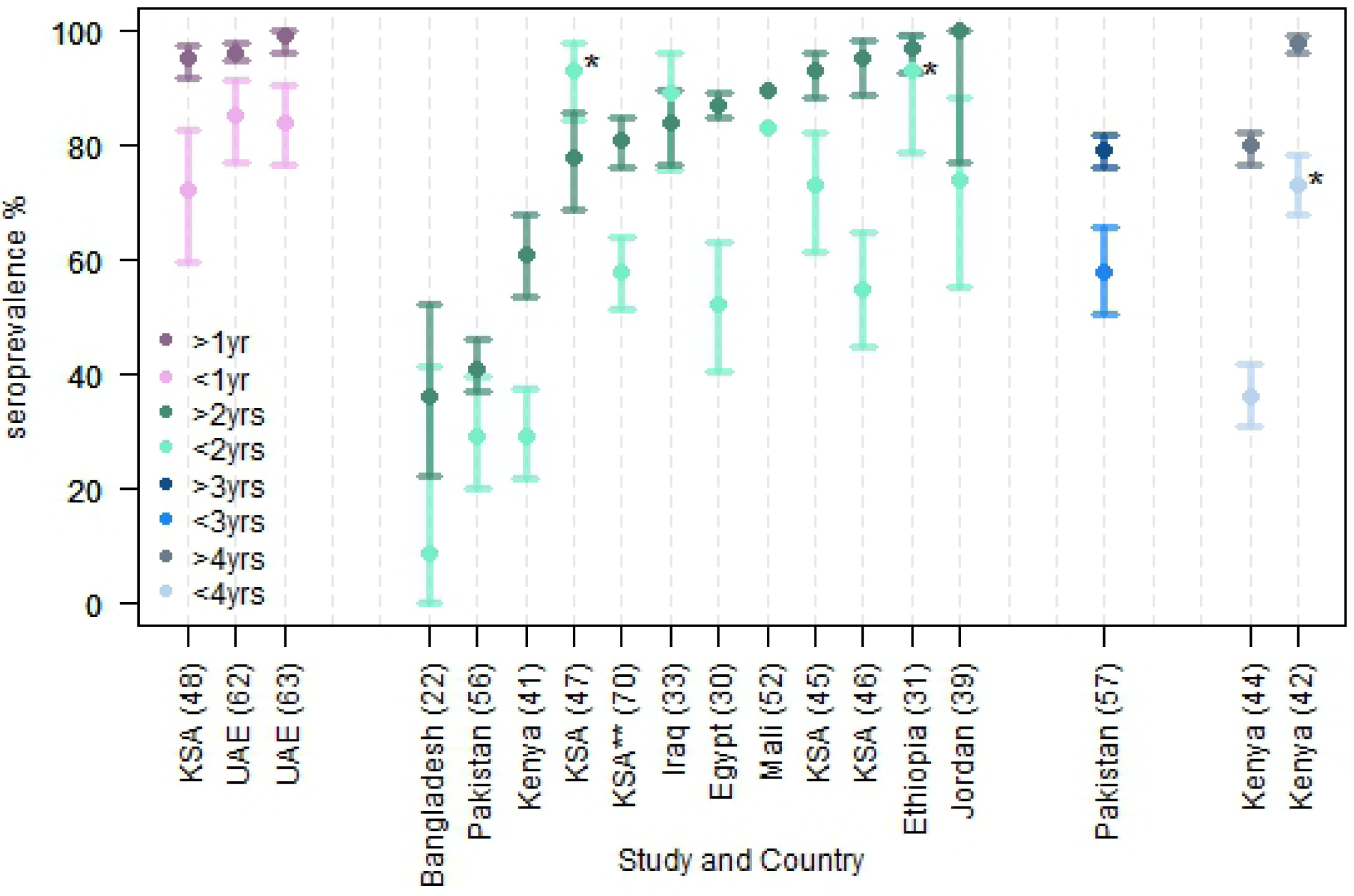
Age stratified seroprevalence measures grouped by available stratification and arranged in order of increasing adult seroprevalence. Bars indicate 95% confidence intervals, calculated by us when not stated in the study, if age class size was available (not available for the population in Mali). *indicates that calves <1-year-old were not included. **indicates that the study was conducted in dromedary populations in response to an epidemiologically linked human MERS-CoV infection.

##### Geographic location

Although MERS-CoV seroprevalence is consistently high across West, North, East Africa and the Middle East at the country level, some studies measured considerable within-country variation in seroprevalence, particularly in Africa (ranges given in Table 1). In Kenya all three studies that presented sub-nationally disaggregated results showed seroprevalence to vary greatly by province. Tunisia, Morocco and Mali also showed considerable regional seroprevalence variation. Less regional variation in seroprevalence was observed within Middle Eastern countries.

##### Sample population characteristics

Several studies found imported animals to have significantly higher seroprevalence than their locally bred counterparts (22, 25, 29, 30). Dromedaries sampled at markets, abattoirs and quarantine sites had higher seroprevalence than those in farms, villages and research facilities (22, 29, 30, 39). In some cases, dromedary origin varied with the type of site sampled suggesting confounding.

In a study conducted across Burkina Faso, Morocco and Ethiopia, dromedaries used for milk and meat had higher seroprevalence than those used for transport (23). In most of the studies that stratified by sex, little difference was seen, but in Kenya females had statistically significantly higher seroprevalence than males (93% vs. 81% in one study (42), and 74% vs. 54% in another (44)) whereas males had significantly higher seroprevalence in Egypt and in KSA (72% and 84% in males vs. 66% in females) (29, 70).

Large and medium herd-size was a significant risk factor for seropositivity across Burkina Faso, Morocco and Ethiopia, as well as nomadic and sedentary husbandry systems as opposed to a mixed lifestyle (23). Conversely, in Kenya there was a non-statistically significant trend for smaller herds to have higher seroprevalence (41). Two other studies in Kenya showed higher seroprevalence in nomadic herds compared with those kept on ranches or those with agro-pastoralist management. However, ranches were in a different region from nomadic herds and the sample size for agro-pastoralist management was very small(40, 42).

##### Prevalence of active MERS-CoV infection – cross-sectional studies

Our search found that dromedary populations in 16 countries have been tested for MERS-CoV RNA, 13 of which report positive results indicating active infection. These include KSA (0.12-56%) (45, 46, 49-51, 70), UAE (0-29% (62–66) or 0-100% if dromedaries epidemiologically linked to human MERS-CoV cases are included(67–69)), Qatar (22-79%) (15, 58, 71, 72), Oman (7%) (55), Iraq (15%) (34), and Jordan (62%) (39), as well as Egypt (4-15%) (27, 29, 30), Ethiopia (10%) (23), Kenya (0.35-0.95%) (43, 44), Nigeria (3-11%) (53, 54), Burkina Faso (5%) (23), Morocco (2%) (23), and Pakistan (3%) (57). See Fig 4 for a map of RNA prevalence, and Table 1). Despite moderate seropositivity, surveys have not detected active MERS-CoV infection in the dromedary populations of Bangladesh or Israel (22, 35).

**Figure 4.**
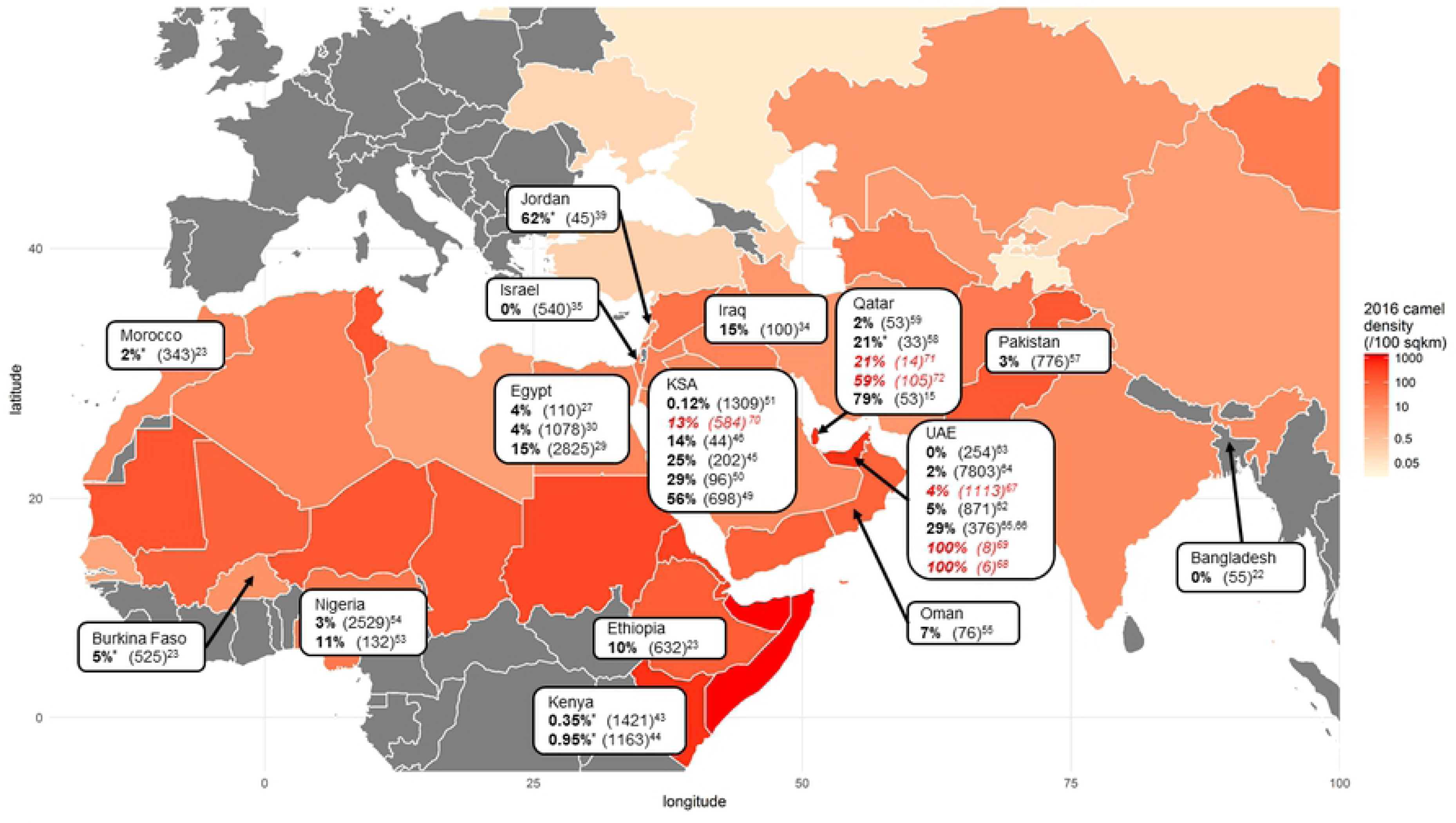
Measures of MERS-CoV RNA prevalence in dromedaries, aggregated at the country level. Total sample size tested is given in parenthesis. Camel density is calculated using FAOSTAT country-level camel population data and World Bank data on country surface area (both for 2016). *value calculated by us from disaggregated sub-national measures of RNA prevalence. Underlined italicised text highlights studies conducted in dromedary populations in response to an epidemiologically linked human MERS-CoV infection.

#### Factors affecting prevalence of infection

##### Age

Age stratified studies in KSA and Jordan found that juveniles had a higher RNA-positivity than adults (39, 45, 70). RNA-positivity also had an inverse association with age in dromedaries across Morocco, Burkina Faso and Ethiopia (23). Two studies in Egypt measured similar positivity rates between juveniles and adults (29, 30).

##### Sample population characteristics

Higher prevalence of RNA shedding was found in imported animals by three studies in Egypt, however site-type is a potential confounder, with local camels being sampled from farms and villages, whilst imported animals were sampled in markets, quarantine centres and abattoirs (27, 29, 30). A study in KSA sampled both local and imported dromedaries within live animal markets found that locally-reared animals had significantly higher prevalence of viral shedding (51). Overall, three studies reported abattoirs and one reported wholesale markets to be associated with an increase in measured prevalence of shedding compared to villages, farms and quarantines (23, 46, 51, 64). Much like seroprevalence, RNA positivity was significantly higher in dromedaries bred for meat or milk compared with those used as transport in Burkina Faso, Morocco and Ethiopia, and shedding was higher amongst females, albeit sex and function were highly correlated (23).

##### Potential temporal trends

Five studies measured RNA prevalence at a defined site at multiple points in time. Animals were not themselves sampled longitudinally. Three studies in Egypt and KSA showed a clear peak in prevalence of viral RNA shedding from December to May (29, 49, 50). The fourth, conducted in wholesale markets in KSA, saw lower rates infection during July and August, and higher positivity in December (51). At an abattoir in Nigeria, no infection was seen from October to mid-January, with prevalence of infection peaking in February after which no more samples were taken (54).

### Evidence of infection and immunity from longitudinal studies

We found 10 longitudinal studies describing 9 incidences of natural infection on farms and in quarantine facilities – 1 in Egypt (29), 4 in KSA (75–78), 5 in UAE (62, 65, 67, 69, 79) and 1 study taking monthly samples of 430 dromedaries in Kenya (43).

### Duration of viral shedding

Four studies of natural infection measured viral shedding in dromedaries at approximately weekly intervals. The maximum time window in which all consecutive nasal samples taken were positive for MERS-CoV RNA ranged from 7-45 days across published studies, with most positive animals becoming negative within 2 weeks (62, 65, 67, 69). All available studies followed animals that were found to be MERS-CoV RNA positive at the first instance of sampling and the duration of shedding prior to sampling is unknown. Further to this, intermittent RNA shedding, and evidence of potential rapid reinfection/coinfection has been observed (65, 67).

### Evidence of Reinfection

Three studies have found dromedaries to be shedding MERS-CoV RNA despite having high antibody titres months or weeks prior to detectable infection. Both older animals whose antibodies reflect past exposure (29, 76, 79), and young calves whose high antibody titres were maternally-acquired immediately post-partum, became infected (79).

One study directly observed recurring infection amongst a herd of dromedaries in Egypt. Four animals were shedding 1-3 months prior to a herd-wide epidemic in which they were RNA-positive once more (29). Sequenced isolates from a market in UAE showed lineage switching from week to week which is also supportive of rapid reinfection or coinfection in calves (65).

Longitudinal studies also indicated that maternally acquired immunity may offer some protection. In both studies of calf-mother-pairs conducted in UAE, MERS-CoV infection became highly prevalent in calves between 4-6 months of age when maternally-acquired antibody titres had waned (62, 79). Samples from reinfected animals have been found to have lower viral loads, suggesting that past infection may ameliorate future infections (76). Viral load and probability of isolating infectious virus were greater when sampling calves, than adults (44, 79).

## Discussion

The results of our systematic review show that MERS-CoV circulates widely in dromedaries across the Middle East and Africa, but transmission varies spatially, and temporally. The sub-national range of MERS-CoV seroprevalence appears to be larger in countries outside of the Arabian Peninsula. Within-country variation in seroprevalence is potentially indicative of differences in transmission dynamics, meaning vaccine strategy evaluation and mathematical modelling will need to be conducted at a sub-national resolution.

The rise of MERS-CoV seroprevalence from 40-90% in juveniles, to 80-100% in adult dromedaries across much of West, North and East Africa and the Middle East, is signature of an endemic disease where the probability of infection increases with time. High seroprevalence in calves <1-year-old in some populations in UAE and KSA suggests high transmission intensity in the Arabian Peninsula, with most dromedaries becoming infected during the first year of life, though maternally acquired antibodies may contribute to seropositivity in young calves(48, 62, 63). The age-dependent seroprevalence values synthesised here should be used to fit models of MERS-CoV transmission in dromedaries and elucidate the likely transmission intensity of the virus in the Middle East and in Africa – a key parameter for estimating vaccination impact. Reporting finer age stratification of young dromedaries would allow a better comparison of transmission intensity in different regions through fitting models of seroconversion.

Despite locally-acquired human cases predominantly being reported within the Arabian Peninsula, the major unilateral trade of camels from the Horn of Africa to the Arabian Peninsula (80) means that the endemicity of MERS-CoV in African dromedary populations has implications for the scope of control programs. Viruses isolated in Africa (Egypt, Ethiopia, Morocco, Nigeria, Burkina Faso and most recently Kenya) have all been classified into Clade C (with West African isolates comprising sub-clade C1(81)), while only Clades A and B have been isolated in the Arabian Peninsula (27, 43, 53, 81). Spike region sequences from Pakistan are similar to those from the Arabian Peninsula (57). Further investigation of the geographical restrictions of MERS-CoV clades would help clarify the extent to which MERS-CoV circulates intercontinentally.

Age-dependent seroprevalence patterns suggest that the higher prevalence of viral shedding in juveniles compared with adults is likely due to immunological naivety. The age-distribution of reported infections synthesised here, suggests that contact with juveniles may pose greater risks of human transmission than adults, making them potential targets for vaccination. However, frequency of human contact with dromedaries may also be animal-age-dependent (62). Calf-focused vaccination may reduce the overall number of dromedary infections but, the reduced risk of exposure would mean that any remaining infections would likely occur at an older age than in the absence of vaccination. It will therefore be important to further investigate the age-dependency of human-dromedary contact patterns and how these vary in different countries and husbandry systems. Vaccination strategies should be evaluated, not only on their likely impact on prevalence of active infection in dromedaries, but also on the age-distribution of infections.

Mapping the movement of dromedaries is necessary to understand the underlying spatial transmission dynamics of MERS-CoV. The mixing of dromedaries underpins interaction between infectious and susceptible individuals and therefore the dynamics of MERS-CoV transmission. Live markets and abattoirs which both had higher prevalence of RNA shedding compared to other site-types in multiple studies, are key locations for animal mixing(23, 28, 51, 64). Quantitative data describing the movement and trading patterns of dromedary populations will be essential for informing models and considering where potential vaccination should take place. A role for markets as drivers of disease dissemination is characteristic of other zoonotic diseases such as avian influenza (82, 83).

Move evidence is required to establish whether MERS-CoV infection in dromedaries is seasonal. The temporal studies in this review observed a peak in prevalence of active infection between December and June (29, 49-51). These were conducted in Egypt and KSA where dromedary calving occurs between October and February (84–86), and Nigeria which has a similar calving season (87). Assuming seasonal calving was driving the trend, and calves become susceptible between 4-6 months (62, 79), we might expect the number of susceptible dromedaries to peak between January and May – which overlaps with the peaks observed. If infection is driven by seasonal calving vaccination would need to occur annually prior to the infection of newly susceptible calves. Based on phylogenetic analysis of MERS-CoV genomes isolated from humans and dromedaries, a seasonal period of elevated risk of zoonotic transmission was estimated to exist from April through to July (6), however, this is not seen consistently in the epidemiology of primary human MERS-CoV cases reported to WHO (88). Further investigation of potential seasonality has been highlighted as a priority by the FAO-OIE-WHO MERS-CoV Technical Working Group (2).

The results of longitudinal studies included in this review demonstrate re-infection of dromedaries despite high titres of MERS-CoV specific antibodies being present in their sera. Unfortunately, the degree and duration of protection afforded by maternally-acquired antibodies and those acquired from infection is unclear. Informative surveys of a better proxy for protective immunity in dromedaries would improve the accuracy of models of reinfection and the likely effects of vaccination.

Although we found maximum duration of RNA shedding to range from 7-45 days across studies, intermittent shedding or rapid reinfection has been seen to occur for 6 weeks which complicates interpretation of RT-PCR derived RNA shedding results for infectious period. A controlled challenge study in 4 dromedaries performed daily sampling and saw the maximum duration of shedding to be 35 days post inoculation, with infectious virus (as determined by plaque assay) isolatable for the first 7 days (89). However, the biological relevance of the challenge dose is not known. More frequent sampling that includes genotyping and captures of the onset of shedding is needed to more accurately estimate the duration of infectiousness following a single natural infection.

Limitations of our study include that a single author completed the systematic search and data extraction. The available studies exhibit differences in study design, criteria for seropositivity, sample-site type and sample population characteristics. Some studies report that the latter two variables are associated with statistically significant differences in seroprevalence or prevalence of infection within individual studies (23, 29, 30, 39, 40, 51, 64). These heterogeneities made quantitative pooling inappropriate.

Available studies do not include camel dense regions of northern Africa or Rajasthan, India, and Yemen. In addition to the countries included in this systematic review of the published literature, OIE has received reports of RNA positive camels in Iran and Kuwait (90, 91). Members at WHO and FAO (Food and Agriculture Organisation of the United Nations) confirm that further RNA testing studies are planned or underway in several countries in Africa (e.g. Ethiopia, Kenya, Egypt, Somalia, Sudan, Algeria and Morocco) and in the Middle East (e.g. Jordan), as well as countries in South East Asia (e.g. Pakistan) (personal communication, Maria D. Van Kerkhove).

## Conclusions

Our findings provide strong evidence that MERS-CoV is endemic in dromedary populations across much of West, North, East Africa and the Middle East, in agreement with the similar systematic review conducted in parallel with our own (11). Calves are likely to play a central role in sustaining circulation of MERS-CoV and should be a target of potential dromedary vaccination. However, the potential for mass vaccination of calves to change the age distribution of infected individuals should be investigated through mathematical modelling of transmission dynamics in dromedary populations and considered in the context of age-dependent human-camel contact frequency patterns. Sites where dromedaries mix may also play a role in driving transmission. A better understanding of dromedary husbandry and trade patterns, as well as quarantine facilities, is needed to identify where dromedaries are infected with MERS-CoV – critical for focussing potential vaccination strategies. Although in a few studies, prevalence of infection appears to peak in the first half of the year, which may be facilitated by the increase in susceptible animals after the calving season, further studies are needed to confirm this. Further longitudinal studies are required to investigate the temporal dynamics of viral shedding and immunity in the animal host and should ideally be capable of distinguishing co-circulating MERS-CoV lineages.

These remaining gaps in our understanding of MERS-CoV transmission dynamics in dromedary populations agree with the prioritized research outlined in the FAO-OIE-WHO Technical Working group report (2) and must be addressed to obtain a clearer picture of what an optimal vaccination strategy would involve, as well as its likely impact, before implementation can be considered further.

## Acknowledgements

None

## Supporting information captions

S1 Appendix: database specific breakdown of the search strategy and search terms used

S2 Appendix: PRISMA flow chart

